# The CLAMP GA-binding transcription factor regulates heat stress-induced transcriptional repression

**DOI:** 10.1101/2023.10.08.561401

**Authors:** Joseph Aguilera, Jingyue Duan, Kaitlyn Cortez, Rachel S. Lee, Angelica Aragon, Mukulika Ray, Erica Larschan

## Abstract

To survive exposure to heat stress (HS), organisms activate stress response genes and repress constitutive gene expression, thereby preventing the accumulation of potentially toxic RNA and protein products. Although many studies have elucidated the mechanisms that drive HS-induced activation of stress response genes across species, little is known about the mechanisms that repress constitutively expressed genes. Using nascent RNA-sequencing, we identify the first reported transcription factor (TF) that regulates repression of constitutive genes upon heat stress across species and define direct and indirect mechanisms of action by integrating 3D genomic approaches. We demonstrate that the CLAMP (Chromatin-linked adaptor for MSL complex proteins) GA-binding transcription factor (TF) regulates ∼75% of the HS-induced repression in *Drosophila,* a well-established model for understanding the mechanisms of HS-induced gene regulation. Using Micro-C, we demonstrate that heat stress induces widespread changes in local 3D chromatin looping, which are significantly associated with HS-induced transcriptional changes. Overall, we identify CLAMP as the first reported TF that induces HS repression, which modulates 3D chromatin looping through both **direct** mechanisms and **indirect** mechanisms. Moreover, we present the highest-resolution heat stress 3D genomic dataset available in *Drosophila,* providing a key resource for generating mechanistic insights into how temperature regulates 3D genomic contacts.

## Introduction

Understanding how external stimuli regulate gene expression is essential for reducing the effects of dynamic changes in abiotic factors on all living organisms. One such external stimulus, temperature, plays a vital role in determining viability and fertility, which often decrease as temperatures rise (1). Heat stress (HS) rapidly changes transcription of both heat stress genes, which become activated, and constitutive genes, which become repressed (2,3). Therefore, understanding the mechanisms that drive changes in transcription upon heat stress will provide critical insight into how transcription rapidly changes in response to stimuli. Because the heat stress response is rapidly inducible, many discoveries of fundamental transcriptional mechanisms have been made using this system, including pausing of RNA Polymerase II, which occurs at thousands of genes across species (4).

Heat stress induces two types of transcriptional changes: 1) activation of heat stress-induced genes; 2) repression of constitutively active genes (5). Best studied in *Drosophila*, decades of research have defined the mechanisms of heat stress-induced gene activation at heat shock-responsive genes, whose promoters contain specific DNA sequences called **H**eat **S**hock Elements (HSEs), bound by Heat Shock Factor (HSF) (4,6,7). In *Drosophila*, the GA-binding transcription factor GAF and transcription elongation factor NELF, which regulates RNA Polymerase II pause release, are also involved in heat stress-regulated gene activation (7–9). Furthermore, in mammals, NELF functions in HS-regulated transcriptional repression mediated by temperature-dependent phase transition (10). However, the mechanisms that target repression specifically to constitutively active genes remain unknown.

In contrast to activation, much less is known about the specific DNA-binding factors that target constitutive genes for repression after heat stress. Transcriptional repression involves a decrease in promoter-proximal paused RNA Polymerase II at repressed genes (3,7) and cryptic premature termination of transcripts with impaired elongation rate and decreased processivity (10) (11). A report also suggests that heat stress induces the relocalization of architectural proteins from TAD borders to Polycomb binding sites at the interior of TADs, which changes 3D chromatin architecture to potentially regulate repression (12). However, the role of 3D chromatin architectural changes during the heat stress response still needs to be better understood, as another report suggests that global 3D architectural changes at the TAD and compartment levels do not occur upon heat stress (2). Neither of these studies was conducted at a sufficiently high resolution to detect specific local 3D chromatin loop changes. Therefore, the mechanisms by which 3D chromatin changes regulate the heat stress response require further study.

Furthermore, HSF and GAF are involved in heat-stress-induced activation but not heat-stress-induced repression (7). GAF has context-dependent synergistic or antagonistic relationships with another GA-binding transcription factor, CLAMP (Chromatin-linked adaptor for MSL complex proteins) (13,14). Although these two transcription factors have similar GA-rich DNA recognition sequences, variation within seemingly similar *cis*-elements drives the context-specific targeting of CLAMP to the male X-chromosome and the histone locus, where it inhibits GAF binding (13,15). Additionally, CLAMP promotes long-range 3D chromatin interactions (16) and interacts with Negative Elongation Factor (NELF), which functions in heat shock repression (10,17). Therefore, we hypothesized that CLAMP regulates heat stress-induced transcriptional repression because it can directly compete for binding with GAF (13), which regulates heat shock activation, and CLAMP interacts with NELF, which regulates repression (17).

To test our hypothesis, we used three approaches: 1) nascent RNA-sequencing (SLAM-seq) to define the function of CLAMP in heat stress-induced transcriptional activation and repression; 2) Micro-C to identify CLAMP-dependent and independent genome-wide, high-resolution local changes in chromatin organization after heat stress, and 3) HiChIP to identify CLAMP-bound 3D chromatin loop anchors associated with heat-stress-dependent transcriptional regulation. We found that, unlike GAF, CLAMP regulates heat-stress-induced transcriptional repression at many more genes than activation, with 75% of all repression being CLAMP-dependent. Moreover, the canonical heat stress-activated genes remain activated in the absence of CLAMP.

Furthermore, we demonstrate that heat stress induces global changes in local 3D chromatin organization at chromatin loop anchors. We found that CLAMP inhibits the formation of 3D chromatin loops during heat stress when it is directly bound to loop anchors, but its occupancy at these anchors does not change. In addition, we compared the occupancy of different insulator proteins and chromatin remodelers at CLAMP-dependent HS-repressed genes prior to heat stress. We found that CLAMP **directly** regulates HS-induced repression through mediating 3D looping when the following factors are enriched near CLAMP binding sites prior to HS: insulator proteins Ibf1/2, ZIPIC, and the chromatin remodeler ISWI, but not Polycomb. In contrast, CLAMP also **indirectly** regulates repression at loci where it is not bound, potentially by either regulating local 3D chromatin loop formation or regulating transcription independent of 3D looping. Future studies are required to identify which factors regulate the indirect effects on 3D looping and transcription in the absence of CLAMP.

Overall, we identify the first DNA-binding factor that directly regulates repression of the majority of genes that are repressed upon heat stress and define how it functions **directly** at sites where it is co-bound with specific cofactors and **indirectly** regulates repression through modulating 3D contacts. Furthermore, we report a key resource to study HS-induced chromatin 3D structural changes because we generate the highest resolution map to date of heat stress-induced 3D genomic contacts. By investigating the mechanism of HS-induced repression, we enriched our overall understanding of how transcriptional repression is regulated in response to environmental cues in a cellular context.

## Materials and methods

### Fly strains and husbandry

*Drosophila melanogaster* fly stocks were maintained at 24°C on standard corn flour sucrose media. Fly strains used: *fkh-GAL4* (Bloomington, #78061), *UAS-CLAMPRNAi[val20]* (Bloomington, #57163). These were crossed to obtain the desired genotype, *fkh-GAL4>UAS-CLAMPRNAi*.

### Cell culture

Kc cells were maintained at 25°C in Schneider’s media supplemented with 10% Fetal Bovine Serum and 1.4X Antibiotic-Antimycotic (Thermofisher Scientific, USA). Cells were passaged every 3 days to maintain an appropriate cell density.

### Polytene chromosome squashes and immunostaining after HS

Third instar larvae from undriven *UAS-CLAMPRNAi* and *fkh-GAL4>UAS-CLAMPRNAi* were incubated for 40 min at 37°C in a water bath inside a microfuge tube plugged with a soft plug (18) and dissected immediately to pull out the salivary gland. Polytene chromosome squashes were prepared using dissected salivary glands as previously described in Reider et al. 2017 (15). We stained polytene chromosomes with rabbit anti-CLAMP (1:1000, SDIX) and mouse anti-RNA pol II (1:500, ab817, Abcam) antibodies. For detection, we used all Alexa Fluor secondary antibodies against rabbit and mouse at a concentration of 1:200 and visualized slides at 40X on a Zeiss Axioimager M1 Epifluorescence upright microscope with the AxioVision version 4.8.2 software.

### Western Blotting to validate *clamp* dsRNA treatment for *clamp* RNAi

Total protein from the samples (1ml of the cell culture): 3 replicates of *clamp* RNAi and GFP RNAi (con) after HS (HS) and without HS (NHS) treated Kc cells for SLAM-seq was extracted by homogenizing cell pellet in the lysis buffer (50 mM Tris-HCl pH 8.0, 150 mM NaCl, 1% SDS, 0.5× protease inhibitor) using a small pestle. The NHS Kc cell samples treated with *clamp* dsRNA and GFP dsRNA for *clamp* RNAi and GFP RNAi used for Micro-C experiment were also used to extract total protein to test Clamp level in *clamp* RNAi vs GFP RNAi Kc cell samples. After a 5 min incubation at room temperature, cleared the samples by centrifuging at room temperature for 10 min at 14,000×*g*. To blot for CLAMP and Actin, 5 µg of total protein was run on a Novex 10% Tris-Glycine precast gel (Life Technologies). Protein was transferred to PVDF membranes using the iBlot transfer system (Thermo Fisher Scientific) and probed the membranes for CLAMP (1:1000, SDIX), Tubulin (1:5000, Abcam), and Actin (1:5000) antibodies using the Western Breeze kit following the manufacturer’s protocol (Thermo Fisher Scientific). Proteins detected on X-ray film (**Figure S1**) using chemifluorescence detection method as mentioned in the kit protocol using AP-tagged secondary antibody and CDP-star substrate provided with the kit (Western Breeze kit, Thermo Fisher scientific, USA).

### SLAM-seq

15 μg each of *clamp* dsRNA and GFP dsRNA were used for *clamp* RNAi and GFP RNAi (con), respectively, per T25 flask. Kc cells were incubated with dsRNA in FBS minus media for 45 minutes and allowed to grow in media supplemented with 10% FBS for 6 days before harvesting. dsRNA targeting *gfp* (control) and *clamp* for RNAi have been previously validated and described (19). PCR products were used as a template to generate dsRNA using the T7 Megascript kit (Ambion, Inc., USA), followed by purification with the Qiagen RNeasy kit (Qiagen, USA).

Two replicates of each for *clamp* RNAi and *gfp* RNAi, HS (Heat stress) samples, and four replicates each for *clamp* RNAi and *gfp* RNAi, NHS (NHS) samples were used for SLAM-seq. Just before harvesting, cells were either incubated in 4SU (4-thiouridine) provided with SLAMseq Anabolic Kinetics Module (Cat. No. 061.24, Lexogen) at RT or at 37°C for one hour in the dark as per the manufacturer’s instructions. Cells were immediately harvested and the pellet was resuspended in 1 mL of Trizol (Invitrogen, USA). RNA isolated in the dark and S4U-labeled transcripts are alkylated with Iodoacteamide as per manufacturer instruction (SLAMseq Explorer and Kinetics Kit user guide, Lexogen). Resulting modified total RNA was used for downstream NGS library preparation using QuantSeq 3‘mRNA-Seq Library Prep Kit for Illumina (FWD) (Cat. No. 015, Lexogen). Libraries sequenced in Illumina Hiseq 4000 with 150x2 bp pair-end sequencing. For analysis Read 2 for the pair-end sequencing was discarded while performing downstream analysis since quality of Read 2 is very low due to the poly(T) stretch at the beginning of Read 2. Data submitted to Gene Expression Omnibus (GSE253645).

### Micro-C

Two replicates each for *clamp RNAi* and *gfp RNAi*, HS (Heat stress) samples and for *clamp RNAi* and *gfp RNAi*, NHS (NHS) samples were used for Micro-C. Kc cells (2X10^6^) were incubated with 15 μg each of *clamp* dsRNA and GFP dsRNA (same as used for SLAM-seq experiment), used for *clamp RNAi* and *gfp RNAi* (con), respectively, per well of a 6-well plate in FBS minus media for 45 minutes and allowed to grow in media supplemented with 10% FBS for 6 days before harvesting. Before harvesting, HS (Heat stress) samples were incubated at 37°C for one hour. 3 × 10^6 cells were used per Micro-C reaction. Cells were aliquoted into 1.5 ml tubes, spun at 500g in a swinging bucket rotor for 5 minutes to pellet the cells, which were placed in -80 °C after removing the supernatant for 1 hour as per the manufacturer’s protocol (Dovetail™ Micro-C Kit user guide version 2.0). Next, cells were thawed at RT, resuspended in 200 μL of 1X PBS (Thermo Fisher Scientific, 10010023), and crosslinked using 0.3M DSG and 37% formaldehyde as per the manufacturer’s protocol (Dovetail™ Micro-C Kit user guide version 2.0). Dovetail™ Micro-C Kit (Catalog # 21006), the DovetailTM Library Module for Illumina (Catalog # 25004), and the primer set for NGS library preparation (DovetailTM Dual Index Primer Set #1 for Illumina, Catalog # 25010) were used for digestion, proximity ligation, library preparation, ligation capture, and amplification. Libraries were subjected to Illumina 150x2 bp pair-end sequencing with 80X coverage. Data submitted to Gene Expression Omnibus (GSE297379).

### HiChIP

Kc Cells were allowed to grow to confluence and harvested. Before harvesting, HS (Heat stress) samples were incubated at 37°C for one hour. 10 × 10^6 cells were used per HiChIP reaction, and two replicates each for NHS (non-heat stress) and HS (Heat stress). Crosslinking was performed using 0.3M DSG and 37% formaldehyde as per the manufacturer’s protocol (Dovetail™ HiChIP MNase Kit user guide). Dovetail™ HiChIP MNase Kit (Catalog # 21007), which includes the Dovetail Library Prep Kit and the primer set for NGS library preparation, was used for digestion, lysate preparation, chromatin immunoprecipitation, proximity ligation, and library preparation. Rabbit anti-CLAMP (5μg/sample, SDIX) was used for the immunoprecipitation step. Libraries were subjected to Illumina 150x2 bp pair-end sequencing. Data submitted to Gene Expression Omnibus (GSE253644).

### Computational analysis

#### a) SLAM-seq Analysis

The nf-core/slamseq (v1.0.0) pipeline was used to process and analyze the SLAM-seq data (20,21). The following parameters were used in the pipeline: -profile singularity, --genome BDGP6, --read_length 150, --trim5 12, --polyA 4, --multimappers true, --quantseq false, --endtoend false, --min_coverage 2, --var_fraction 0.2, --conversions 1, --base_quality 27, --pvalue 0.05, --skip_trimming false, --skip_deseq2 false. Adapter trimming was performed using Trim Galore (0.6.5). Conversion-aware mapping, alignment filtering, multimapper recovery, SNP calling, read quantification, gene-level quantification collapsing, QC stats, and result summarizations were all performed using SlamDunk (0.4.3)(20). Differentially expressed genes were then determined using DESeq2 (1.22.1).

#### b) Micro-C analysis

Micro-C raw data was analyzed using the HiC-Pro toolkit (3.1.0) with default parameters. Normalized contact maps deriving from HiC-Pro were converted to the cooler format using utility scripts (https://github.com/nservant/HiC-Pro/tree/master/bin/utils) in the HiC-Pro toolkit (3.1.0). Subsequently, the cooler contact maps for each condition were used to identify differential chromatin interactions using the HiCcompare library (1.24.0) in Bioconductor (3.18). Significant differential chromatin interactions (P < 0.01) were used in a range of analyses throughout the study. Differential loops within our statistical cutoff were visualized in a MA plot in Python (3.8) using matplotlib (3.7.5) and seaborn (0.13.2). Furthermore, the bedtools suite (2.31.0) was used to perform overlaps (bedtools pairtobed) with dm6 genes, determining the number of genes that were localized within loops and their anchors, respectively. The Mann-Whitney test was used to determine the statistical significance of differential loop sizes between our studied conditions. Moreover, differential loops were visualized and compared with other datasets in the study using the coolbox API (0.3.9) (22).

#### c) HiChIP Analysis

HiChIP raw data was analyzed using the Dovetail genomics Hi-ChIP documentation (https://hichip.readthedocs.io/en/latest/index.html). Sequences were mapped to the dm6 genome using the Burrows-Wheeler Aligner (0.7.17) (23) passing the following arguments: mem, -5, -S, - P, -T0; default values were used for all other settings. Using the parse module of pairtools (0.3.0), valid ligation events were recorded; the following parameters were changed from default values: --min-mapq 40, --walks-policy 5unique, --max-inter-align-gap 30. PCR duplicates were removed using the dedup module of pairtools (0.3.0). Subsequently, the split module of pairtools was used to generate the final bam and pair files. The samtools (1.17) sort module was used to sort the bam files. To determine the quality of the proximity ligation events, scripts generated by the Dovetail Genomics team were utilized, which are deposited in the following GitHub repository (https://github.com/dovetail-genomics/HiChiP).

The cooler suite (0.9.3) (24) was used to generate contact maps from pair files, and the coolbox API (0.3.9) (22) was used to visualize the contact maps and bigwig tracks. To call loops, the flexible FitHiChIP tool (10.0) (25) was used as resolutions are easily adjustable, and it can detect different categories of 3D chromatin interactions. Furthermore, since FitHiChIP required 1D peaks from the same tissue and conditions, MACS2 (2.2.9.1) was used to call peaks on the Hi-ChIP data. Differential loops between conditions were determined using edgeR in custom scripts located in the FitHiChIP repository (https://github.com/ay-lab/FitHiChIP) and were visualized in a MA plot in Python (3.8) using matplotlib (3.7.5) and seaborn (0.13.2). Differential loops were visualized using APA plots with the coolpup.py (61) suite (1.0.0), displaying average signal over loop focal points. The central pixel within the APA plot is the central bin (2.5 kb), which contains the differential chromatin contact. The total size of the region displayed, including the central bin, is represented within the figures legends or within plot. Scores represented in the top left corners of the APA plot correspond to the central pixel intensity relative to the background. Furthermore, the bedtools suite (2.31.0) was used to perform overlaps (bedtools pairtobed) with dm6 genes, determining the number of genes that were localized within loops and their anchors, respectively. The Mann-Whitney test was used to determine the statistical significance of differential loop sizes between our studied conditions.

#### d) Integration of Micro-C, HiChIP, and SLAM-seq datasets

To overlap Micro-C and HiChIP (3D genomics) with SLAM-seq (nascent RNA sequencing), we first had to transform the output of each technique into a mutually comparable format. Therefore, to analyze all three datasets, Micro-C and HiChIP differential loops were annotated with FlyBase genes, and SLAM-seq differential transcripts were converted to FlyBase genes. Micro-C and HiChIP differential loop anchors were annotated using a custom Python script and the bedtools suite (2.31.0). In addition, the entire loop span was annotated using the bedtools suite (2.31.0), collecting all FlyBase genes that were localized at the loop anchors and within the loop. Using this method, we derived gene lists from the differential loop analyses of Micro-C and HiChIP, which we could compare with the SLAM-seq differential gene lists. All overlaps presented were visualized using the matplotlib-venn (0.11.9) Python package, and Fisher’s Exact test combined with FDR correction was used to determine statistical significance between all gene lists (**Table S1**). The coolbox API (0.3.9) (22) was used to visually compare all datasets in the study, observing differential contact maps, chromatin loops, ChIP signal, nascent RNA signal, and genes within a specified region.

#### e) ChIP-seq Analysis

Raw data (GSE118047, GSE30740, GSE36374, GSE36393, GSE54529, GSE63518, GSE80702, GSE89244) (12,26–33) was filtered using Trim Galore (0.5.0), then aligned to the dm6 genome using the mem module of the Burrows-Wheeler Aligner (0.7.17) with default settings. BigWig tracks were produced using the bamCoverage module in the deeptools suite (3.2.1) with the following parameters: --binSize 50, --normalizeUsing CPM, --extendReads 200, --exactScaling, --centerReads, --blackListFileName dm6-blacklist.bed. The respective BigWig tracks were visualized using the coolbox API (0.3.9) and the deeptools suite (3.2.1) over regions of interest.

## Results

### 1. CLAMP is required for the repression of transcription in response to heat stress

To understand how the CLAMP GA-binding transcription factor regulates nascent transcription after heat stress, we performed high-throughput sequencing of nascent RNA in *Drosophila* Kc cells using Thiol (SH)–linked alkylation for the metabolic sequencing of RNA (SLAM-seq) (34). SLAM-seq quantifies nascent mRNA by measuring 4SU (4-thiouridine) incorporation as thymine-to-cytosine (T to C) conversions. Cells were incubated in 4SU either at room temperature (RT) for one hour (No heat stress, NHS) or at 37°C for one hour (Heat stress, HS), and then immediately processed to identify which mRNAs incorporated 4SU within the total mRNA pool. Next, we compared the mRNAs incorporating 4SU under HS and NHS conditions to determine which transcripts had increased levels after HS (activated genes) and which transcripts had reduced levels after HS (repressed genes) (padj ≤ 0.05 and fold change ≥ 1.5).

Consistent with a prior report (7), we identified a subset of transcripts that were activated (**N = 995**) and a larger subset that was repressed (**N = 3,547**) in control (*gfp RNAi)* cells after heat stress (HS) (**Figure 1A**). HS-activated transcripts encompass all known heat stress-responsive genes, including those for heat shock chaperone proteins (**Table S2a**). Much less is known about the mechanisms that mediate HS-induced repression compared to activation. We hypothesized that HS-induced repression is mediated by the CLAMP GA-binding protein TF because it is known to compete with GAF, which regulates heat-stress-induced activation (7,13). Since CLAMP also regulates male X-chromosome dosage compensation in addition to global transcription in both males and females, we decided to examine the function of CLAMP during the heat stress response in the Kc female cell line, which does not undergo dosage compensation.

**Figure 1.**
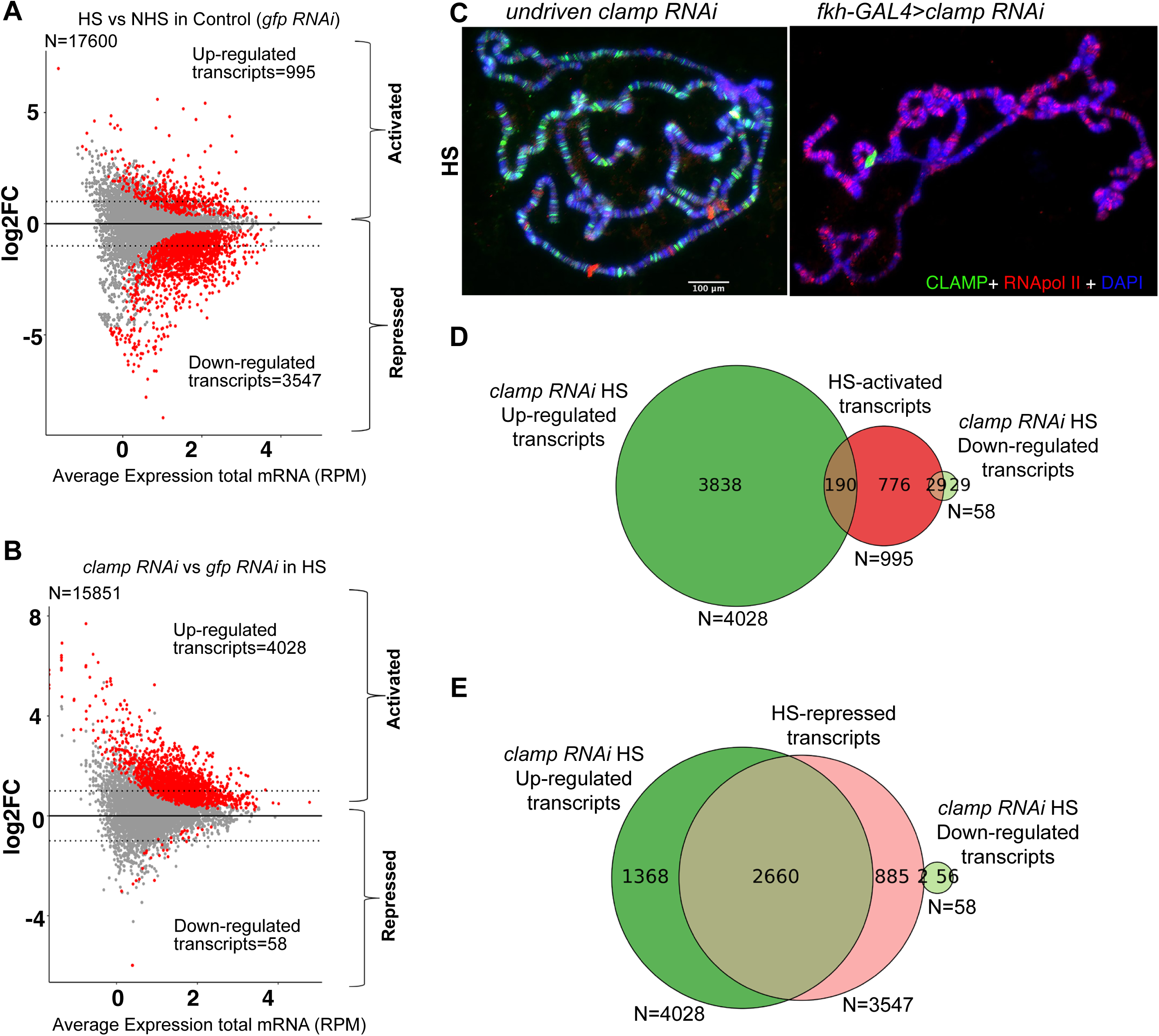
CLAMP regulates heat-stress-induced transcriptional repression more strongly than activation. **A-B.** MA plots showing significant changes (FC ≥ 1.5 and padj ≤ 0.05) in the levels of nascent transcripts (red dots) after HS (heat stress) compared to the control NHS (No heat stress) condition in control (*gfp RNAi*) cells (**A**) and after HS (heat stress) in the absence of CLAMP (*clamp RNAi*) compared to control (*gfp RNAi*) cells (**B**). Dotted horizontal lines in **A** and **B** denote ±1 log_2_ FC change cutoffs. **C.** Immunofluorescence images of HS (37⁰C for 1 Hr) salivary gland polytene spreads showing the distribution of active RNA polymerase II (red) on chromatin (DAPI: blue) in the presence (undriven *clamp RNAi*) and absence (*fkh-GAL4>clamp RNAi*) of CLAMP (green). The scale bar (100 μM) applies to both panels in **C**. **D-E.** Venn diagrams comparing HS-activated transcripts (deep red circle in **D**) and HS-repressed transcripts (light red circle in **E**) from control cells (*gfp RNAi*) to *clamp RNAi* HS Up-regulated (dark green circle) and *clamp RNAi* HS Down-regulated (light green circle) transcripts in *clamp* RNAi-treated cells.

In the absence of heat stress (NHS), our validated *clamp* RNAi treatment (**Figure S1**, 13,35,36) causes increased levels of 987 nascent transcripts and decreased levels of 246 nascent transcripts (**Figure S2A**). Therefore, CLAMP functions as a repressor more frequently than an activator in Kc cells before heat stress (**Figure S2A, Table S2b**). After HS, unlike control (*gfp RNAi)* cells (**Figure 1A, Table S2a**), in *clamp* RNAi*-*treated cells, we observed very little down-regulation of ongoing transcription (**Figure S2B, Table S2c**). Furthermore, we observed a widespread increase in transcript levels in *clamp* RNAi*-*treated cells compared with control *gfp* RNAi*-*treated cells (**4,028** upregulated transcripts; **58** downregulated transcripts) (**Figure 1B, Table S2d**). Therefore, CLAMP functions more frequently as a repressor than an activator of transcription after heat stress in cell culture.

Next, we tested whether CLAMP promotes repression of heat stress response genes *in vivo*. We examined the distribution of active RNA Polymerase II (Pol II) on *Drosophila* larval salivary gland polytene chromosomes after HS (37°C) in control (undriven *clamp RNAi*) and salivary gland driver (*fkh-GAL4*>*clamp RNAi*) larvae (**Figure 1C**). Consistent with extensive prior work (18,37), heat stress restricts the binding of active Pol II (red) to specific cytological positions corresponding to HS-activated loci on control polytene chromosomes (**Figure 1C**). In contrast, after *clamp* RNAi, active Pol II (red) remains bound genome-wide, including at essential HS-activated loci, suggesting a loss of HS-induced repression but not activation. Notably, *clamp RNAi* flies retain CLAMP at the histone locus, consistent with prior reports (**Figure 1C**), perhaps because it contains 200 CLAMP binding sites in close proximity to each other (15). Overall, our data show that CLAMP is a stronger regulator of heat stress-induced repression than activation in both Kc cells and flies.

To determine whether CLAMP regulates HS-induced gene activation, we measured the effect of *clamp* RNAi on transcripts that were activated after HS in control cells (**Figure 1D**, **Table S2a, c**). We found that only 2.9 % (29 out of 995) of transcripts activated in control *gfp* RNAi cells are dependent on CLAMP (**Figure 1D**). In contrast, CLAMP inhibits further activation of 19% (190 out of 995) of HS-activated transcripts (**Figure 1D**). Furthermore, all well-studied canonical HS-induced transcripts, including the *hsp70* gene (7), are still induced even after *clamp* RNAi treatment (**Table S2d**).

Next, we defined the set of transcripts and genes that require CLAMP for repression after HS (CLAMP-dependent HS repression) by comparing HS-repressed transcripts (N=3547) with transcripts specifically up-regulated in *clamp RNAi*-treated cells (N=4028) and not in control cells (*gfp RNAi*) both under HS conditions (**Figure 1E**). We found that 75% (2660/3547) of transcripts are CLAMP-dependently repressed after HS (**Figure 1E, Table S3**). At the gene level, we identified HS-activated (N=421) and HS-repressed (N=1047) genes (**Figure S3A, B**), and found that in total 1061 genes are up-regulated and only 30 are down-regulated after HS in the absence of CLAMP (*clamp RNAi*). There is a significant overlap between HS-repressed genes and *clamp RNAi* HS upregulated genes (**Figure S3B,** N=735, p=0.00). All overlaps presented were subjected to Fisher’s Exact test combined with FDR correction to determine statistical significance between comparisons (**Table S1**). Thus, the loss of CLAMP has a more substantial effect on HS-induced repression (75% of repressed transcripts are CLAMP-dependent, **Table S3**) than on activation (2.9% of activated transcripts are CLAMP-dependent, **Table S3**).

### 2. Heat stress induces widespread local 3D chromatin changes at chromatin loop anchors

We next asked the question: How does CLAMP regulate transcriptional repression upon HS? Since heat stress induces aggregation of proteins and RNA, especially newly synthesized protein molecules (38), we predict that repression of ongoing transcription ensures that during heat stress, the process of protein production is inhibited at the transcriptional level. Thus, in order to survive under stress by conserving energy and preventing the formation of potentially toxic protein aggregates, the mechanism regulating repression of transcription upon heat shock (HS) needs to be essentially a global phenomenon. Therefore, we hypothesized that HS-induced global changes in transcription repression may be regulated by genome-wide changes, such as global changes in 3D chromatin architecture. Additionally, CLAMP is known to regulate 3D chromatin loop formation and global changes in chromatin organization (16), which can be sensitive to changes in temperature, likely through phase transition (39), which CLAMP can undergo (40).

The function of 3D chromatin organization in the response to heat shock (HS) has remained unclear because the resolution of prior methods was low, and newer high-resolution techniques were not available when these experiments were conducted. One prior report suggests that HS induces the re-localization of architectural proteins from TAD borders to increase long-range interactions enriched with Polycomb marks, thereby regulating repression (12). In plants, HS alters the 3D arrangement of promoter-enhancer interactions to regulate the activation of heat shock-responsive genes (39,41,42). In contrast, other reports could not find a correlation between global three-dimensional chromatin changes and transcriptional changes after heat shock (2). Unfortunately, prior studies on 3D chromatin and HS used the Hi-C technique with a resolution of ∼40 kb at anchor regions, making it challenging to understand the function of specific chromatin-associated factors that bind to regions that are approximately 0.1 kb (61). However, using the newer micro-C technique to measure 3D chromatin changes after HS, at a resolution of 1-5 kb, could make it possible to define local changes in chromatin organization marked by chromatin loops (43) that are regulated by a TF like CLAMP.

Thus, using the micro-C technique, we first investigated global changes in 3D chromatin loops at a resolution of 2.5kb after HS in control samples (*gfp RNAi*) (**Figure 2A**) and then HS induced global changes in chromatin loops regulated by CLAMP (**Figure 2B**) by performing the micro-C method in *clamp* RNAi-treated cells under HS and NHS conditions using a resolution of 2.5 kb to examine gene-level transcriptional changes (**Figure 2C**). Moreover, this is the first high-resolution *Drosophila* data set, identifying 3D chromatin changes associated with heat stress that others can mine for 3D chromatin changes associated with HS at specific genomic sites.

**Figure 2.**
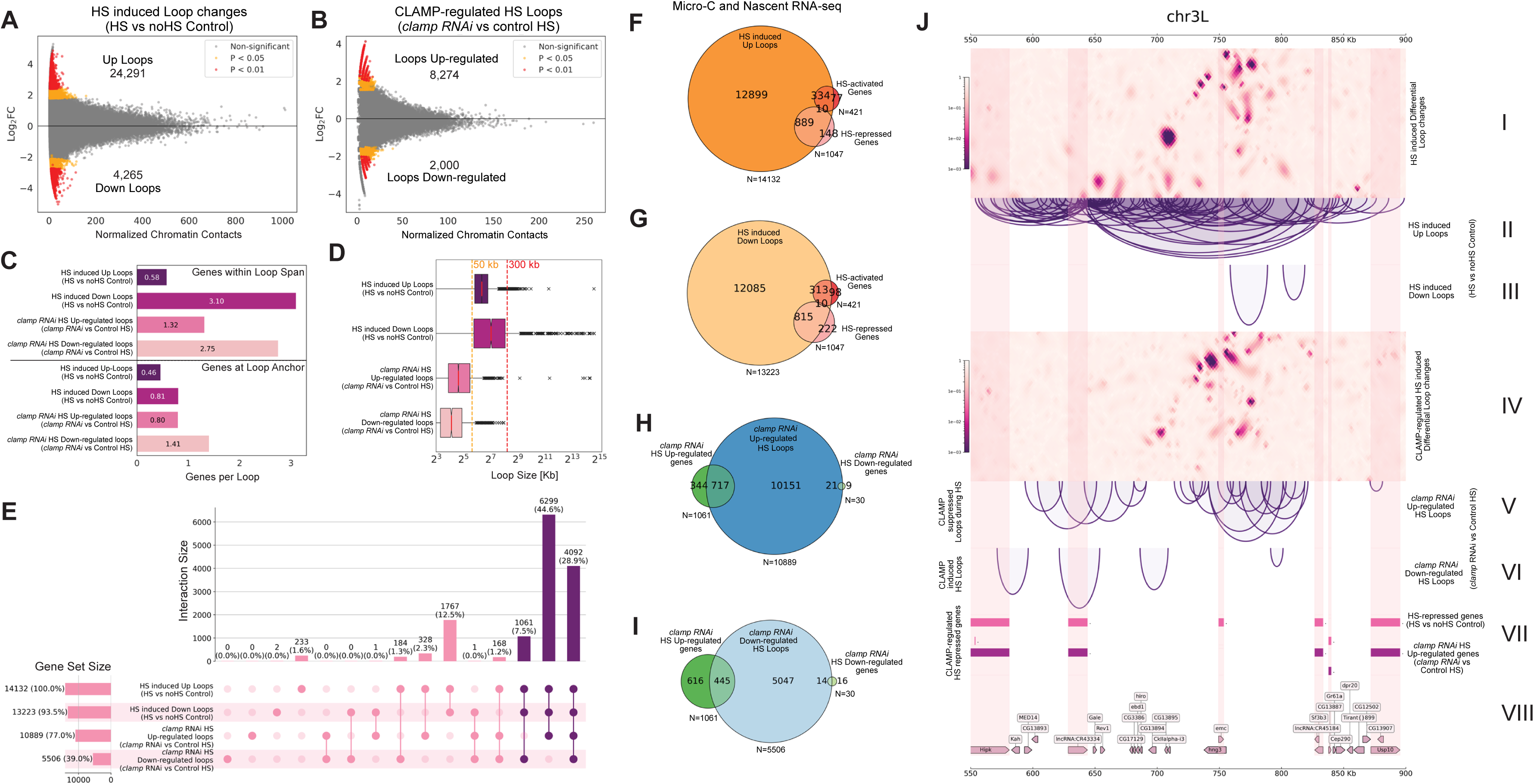
Heat stress induces widespread local 3D chromatin changes at the loop anchors. **A-B.** MA plots for **differential** loop anchors gained upon HS (HS induced Up loop anchors) or loop anchors lost upon HS (HS induced Down loop anchors) in control cells (**A,** Up loops**=**24,291, Down Loops=4,265) and *clamp RNAi* cells (**B,** Loops Up-regulated**=**8,274, Loops Down-regulated =2,000). p<0.01 **C.** Bar plots show the average number of genes associated with (top box) and present at (bottom box) the HS-induced genome-wide chromatin loop anchors and HS-induced CLAMP-dependent chromatin loop anchors. **D.** Box plots showing average loop size of HS-induced genome-wide changing chromatin loops and CLAMP-dependent HS-induced differential chromatin loops with significant difference between average chromatin HS-induced up and down loop sizes, both in case of HS-induced genome-wide changing chromatin loops (Mann–Whitney Test, FDR corrected p=5.63E-186) and CLAMP-dependent HS-induced differential chromatin loops (Mann–Whitney Test, FDR corrected p=3.97E-70), and also between HS-induced genome-wide changing and CLAMP-dependent HS-induced differential chromatin loops themselves (Mann–Whitney Test, FDR corrected p=0) **E.** Upset Plot showing genes associated with differential chromatin loop changes and cardinality of every category combination seen as chromatin loop changes after HS in control (top two rows) and *clamp* RNAi-treated (bottom two rows) cells. The first four columns show chromatin loop-associated genes only in each category, and the following six columns show those in each combination of exactly two named sets, the four columns after that combination of three named sets, and the last column shows genes associated with chromatin loop changes in all four categories. HS-associated CLAMP-dependent chromatin loop changes are mostly represented in the last three columns/bars. **F-I.** Venn diagrams comparing HS-associated chromatin loop changes (genes within the loop) and HS-activated and repressed genes (**F** and **G**); as well as CLAMP-dependent chromatin loop changes during HS with CLAMP-dependent HS-induced transcriptional changes (**H** and **I**), showing significant (Fisher’s Test) overlap between HS-associated Up loops (**F**) and Down loops (**G**) with HS-repressed genes; and CLAMP-dependent HS-associated Down loops (**H,** *clamp RNAi* up-regulated HS-loops) and Up loops (**I,** *clamp RNAi* down-regulated HS-loops) with CLAMP-dependent HS-repressed genes (*clamp RNAi HS* up-regulated genes). FDR corrected **p-values** are 7.487 E-09 (**F**), 2.53 E-4 (**G**), 2.0 E-6 (**H**), and 4.290 E-15 (**I**), respectively. **J.** 350 kb region on 3L showing differential contact interaction results of selfish algorithm on HS and NHS micro-C contact map (**I**) with associated chromatin loops that are formed (**II**) and inhibited after HS (**III**). CLAMP-dependent differential contact interaction results of the selfish algorithm are shown on *clamp* RNAi-treated HS and control HS micro-C contact maps (**IV**) with CLAMP inhibited (**V**) and favoured (**VI**) chromatin loops. The color bar in **I** and **IV** indicates the q-value (BH adjusted *p-*value) produced from the DCI analysis. Darker color means this interaction has a lower q-value; that is to say, two contact maps are more diverse at this location. Bed files for differential nascent RNA-seq (SLAM-seq) data from HS and NHS *clamp* RNAi-treated and control cells show the CLAMP-dependent HS-repressed genes (**VII**) among the corresponding genes within the genomic region associated with the loops (**VIII**)

First, we identified widespread HS-induced changes in chromatin loops that were increased in frequency (Up loops, N = 24,291) and those decreased in frequency (Down loops, N = 4,265) by HS (**Figure 2A, p<0.01**). Interestingly, *clamp* RNAi enhanced HS-induced loop formation (*clamp* RNAi HS Up-regulated Loops, N = 8,274) more frequently than it decreased HS-induced loop formation (*clamp* RNAi HS Down-regulated Loops, N = 2,000). Thus, during heat stress, CLAMP primarily functions as an inhibitor of chromatin loop formation (**Figure 2B, p<0.01**).

Next, we defined how genome-wide local 3D changes in chromatin loop formation correspond to gene distribution at two classes of sites: 1) within the chromatin loops and 2) at the loop anchors (**Figure 2C**). This analysis allowed us to determine if HS-associated chromatin loop changes are related to HS-induced transcriptional repression by comparing the chromatin-loop-associated genes with CLAMP-dependent HS-repressed genes (**Table S4**). As expected, at the loop anchors which are 2.5 kb in length approximately one gene was associated within both HS-induced Up loops and Down loops independent of *clamp* RNAi, because the average gene size is 3 kb in *Drosophila* (**Table S5**). However, when we evaluated the number of genes within the loops, we found significant differences across subcategories with down-regulated loops spanning more genes (3 genes) than upregulated loops ( 1 gene) independent of *clamp RNAi*. (**Figure 2C, Table S6**).

Next, we tested whether there were more genes in a loop because the loops were larger in length or because the genes were more densely packed within the same sized loop (**Figure 2D, Table S1**). For HS-induced chromatin loops, the HS-induced Down loops were significantly larger (avg 627 kb) than HS-induced Up loops (avg 95 kb), and thus had more genes within each loop (**Figure 2C, D**). However, interestingly, the average size of both *clamp* RNAi HS Up-regulated and Down-regulated loops was significantly lower than that of control HS-induced chromatin loops. Therefore, CLAMP regulates the chromatin contacts that form shorter gene-rich loops. Moreover, CLAMP regulates the majority (∼85%) of all HS-associated changes in chromatin loops, and most of the time it prevents their formation (**Figure 2E**).

After determining that CLAMP regulates HS-induced chromatin changes at gene-rich genomic regions, we asked whether these HS-associated 3D chromatin changes were related to HS-induced transcriptional changes (**Figure 2F-I**). We compared HS-regulated genes with genes located between (**Figures 2F-I**) and at loop anchors (**Figures S4A-D**) of the HS-induced chromatin loops. Almost all HS-activated and HS-repressed genes overlapped with HS-induced loop changes (**Figures 2F and 2G, Table S1**). This was unsurprising considering we identified whole-genome-wide chromatin loop changes induced by HS. However, we also found that most CLAMP-dependent HS-repressed genes (*clamp RNAi* HS Up-regulated genes) overlapped with *clamp* RNAi HS Up-regulated and Down-regulated loops (**Figures 2H and 2I, Table S1**). Interestingly, more (**N=717, 68%**) of the CLAMP-dependent HS-repressed genes overlapped with HS loops suppressed by CLAMP (*clamp* RNAi HS Up-regulated loops) compared with HS loops promoted by CLAMP (**N=445, 42%**) (**Figures 2H-I).** Overall, these data are consistent with a function for CLAMP in repressing heat shock-induced gene expression by reducing the formation of chromatin loops that normally activate genes.

To visualize the association of CLAMP-mediated chromatin loop changes with CLAMP-dependent HS-repression, we identified a 350 kb gene-dense region on chromosome 3L that is associated with a large number of HS-induced chromatin loops and multiple HS-repressed genes (**Figure 2J**). CLAMP regulates a subset of the chromatin loops in this region which are associated with CLAMP-dependent HS-repressed genes. Thus, we demonstrate that CLAMP regulates 3D chromatin changes during heat shock (HS) which are associated with CLAMP-dependent HS-induced transcriptional repression.

### 3. CLAMP inhibits chromatin loop formation during HS and regulates repression of associated genes through both direct and indirect mechanisms

The next question we asked was whether CLAMP has a direct effect on 3D chromatin changes and related transcriptional changes after HS or if it functions through a combination of direct effects at CLAMP-bound loci and indirect mechanisms as is common for most TFs. High-resolution chromatin organization data in HS *clamp* RNAi cells reveal that CLAMP regulates chromatin loop rearrangement during HS (**Figure 2B**) at loops associated with CLAMP-dependent repressed genes (**Figure 2I, J**). However, to understand the direct role of CLAMP in transcriptional regulation after HS at high resolution in 3D, we used HiChIP to define which changes were directly associated with CLAMP binding to loop anchors. HiChIP measures TF binding and 3D loop anchors simultaneously at less than 1kb resolution (25,26,44), making it possible to define both TF binding and associated 3D chromatin loops. Therefore, we used HiChIP to investigate whether changes in 3D loop anchor contacts are associated with alterations in CLAMP occupancy on chromatin and HS-induced transcriptional repression.

We performed HiChIP using validated antibodies (15–17) targeting the CLAMP protein to identify CLAMP-associated loop anchors in Kc cells exposed to HS and matched control NHS Kc cells. Loop anchors bring distal regulatory elements closer to each other, which can regulate activation or repression of gene expression depending on the factors bound (26,45). Using the FitHiCHIP platform (25), we first identified the following categories of CLAMP-bound chromatin loops (**Figure S5**): 1) present in control cells without HS (NHS) (**N=1515**); 2) shared loops that are present in both control (NHS) cells and HS cells (**N=421**); 3) loops present only after HS and not in NHS conditions (**N=563**). We then classified loops into subcategories based on whether they <u>increased</u> or <u>decreased</u> after heat shock: 1) CLAMP-bound HS induced Up loops gained, and 2) CLAMP-bound HS induced Down loops lost upon HS (**Figure 3A, B**). We found that most CLAMP-bound chromatin loops are lost (**N=980, Down loops**) upon HS and very few are gained (**N=71, Up Loops**), clearly showing that CLAMP-bound chromatin loops are mainly down-regulated after HS. Our analysis shows that of the 980 CLAMP-associated HS induced down loops (HiChIP), 931 loops (95%) only had one anchor associated with CLAMP direct binding, while 49 loops (5%) had both anchors associated with CLAMP direct binding. Of the 71 CLAMP-associated HS induced up loops (HiChIP), all 71 loops (100%) only had one anchor associated with CLAMP direct binding. Overall, we conclude that the **direct** effect of CLAMP is to promote the loss of 3D chromatin loops at specific genomic locations after HS more frequently than it promotes the formation of new loops.

**Figure 3.**
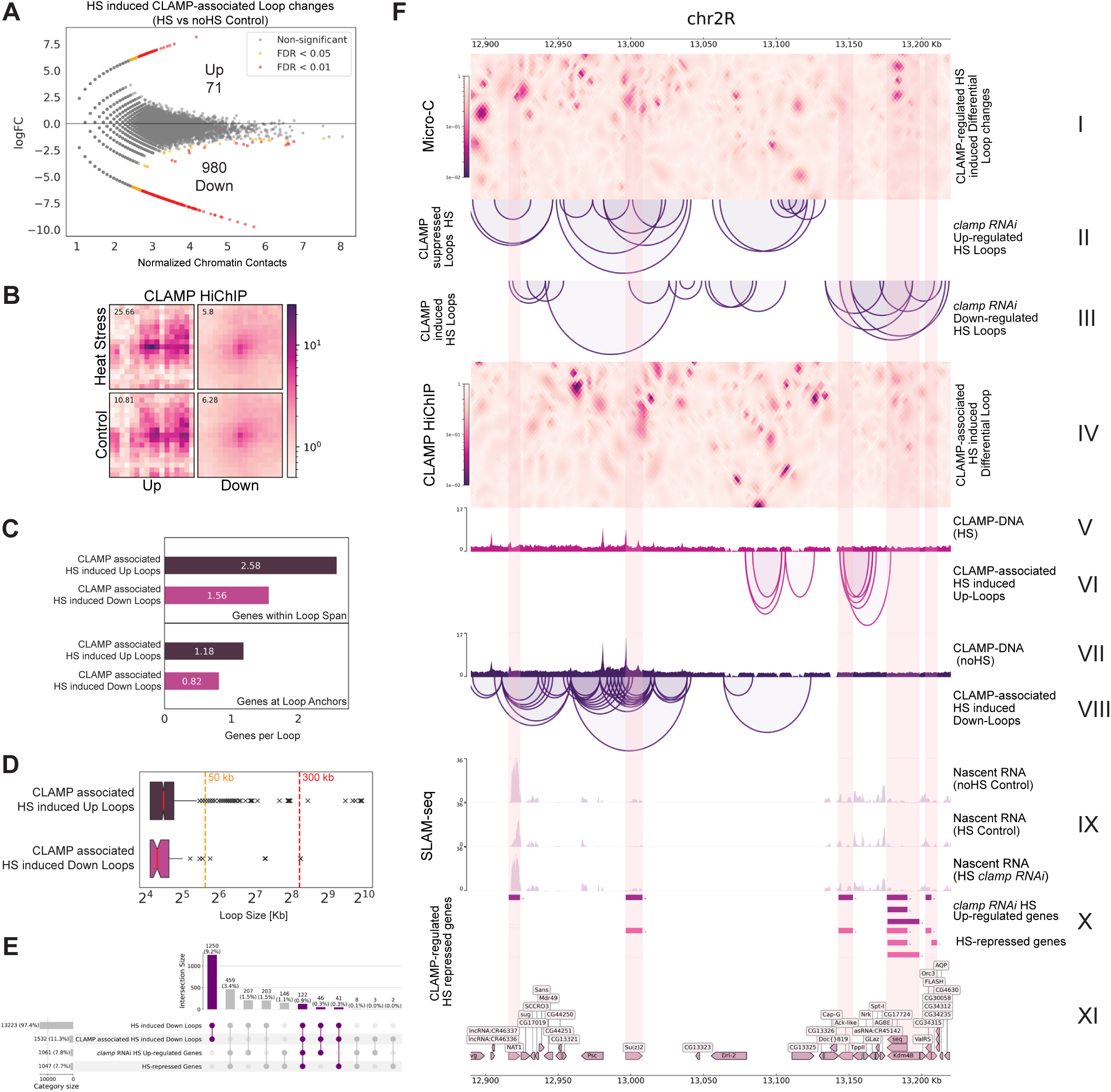
CLAMP regulates chromatin loop changes during HS, directly as well as indirectly, to repress gene transcription. **A.** 2D metaplot of CLAMP HiChIP data centered on significant interactions identified from CLAMP HiChIP under NHS or HS conditions for **differential** loop anchors gained upon HS (HS Up loop anchors, N=71) or loop anchors lost upon HS (HS Down loop anchors, N=980). The score indicates the enrichment of the center pixel compared with the top left corner. **B.** CLAMP-bound loop pile-ups for HiChIP data from heat stress (HS) and control cells showing loss and gain of chromatin structures marked by chromatin loops defined in HS and control cells. The color is average interaction frequency across the entire dataset surrounding the defined HS Up and Down loops. A high signal indicates strong interaction between loop anchors. ±20 kb padding was applied to the central 2.5 kb bin, displaying a 42.5 kb aggregated region. **C.** Barplots showing average number of genes associated within (top box) and present at (bottom box) the CLAMP-bound chromatin loop anchors up and down-regulated after Heat shock (HS) **D.** Box plots showing significant difference (Mann–Whitney Test, FDR corrected p=0.0354) between average loop size of CLAMP-bound chromatin loop anchors up and down-regulated after Heat shock (HS). **E.** Upset Plot comparing all (Micro-C data) and CLAMP-bound (HiChIP data) chromatin loops lost after HS with HS-repressed and CLAMP-dependent HS-repressed genes, showing significant (Fisher’s Test) overlap between CLAMP-bound HS-associated Down-loops with global genome-wide loss of HS-associated loops (p=1.26E-115), HS-repressed genes (p=9.59E-18), and CLAMP-dependent HS-repressed genes (*clamp RNAi HS* up-regulated genes, p=2.57E-19). **F.** 300 kb region on 2R showing CLAMP-regulated differential contact interaction results of selfish algorithm on *clamp* RNAi-treated and control HS micro-C contact map (**I**) with associated CLAMP suppressed (**II**) and induced (**III**) chromatin loops after HS. CLAMP-associated differential contact interaction results of the selfish algorithm are shown on HS and NHS CLAMP HiChIP contact maps (**IV**) with CLAMP-bound (**V** and **VII**) directly associated Up (**VI**) and Down (**VIII**) chromatin loops. The color bar in **I** and **IV** indicates the q-value (BH adjusted *p*-value) produced from the DCI analysis. A darker color means this interaction has a lower q-value; that is to say, two contact maps are more diverse at this location. Normalized reads and bed files for differential nascent RNA-seq (SLAM-seq) data from HS and NHS *clamp* RNAi-treated and control cells show the CLAMP-dependent HS-repressed genes (**IX and X**) among the corresponding genes within the genomic region associated with the loops (**XI)**

Next, we asked whether CLAMP occupancy on chromatin changes when 3D loop contacts change. Therefore, we categorized CLAMP-bound loop anchors lost after HS (HS Down) and gained after HS (HS Up) into the following sub-classes depending on the change in occupancy of CLAMP on chromatin (similar to ChIP-seq) which HiChIP also quantifies in parallel with 3D loop anchors (**Figure S6**): 1) **H**igh **D**ifference (**HD**) when CLAMP occupancy between HS and NHS conditions is differential by EdgeR (46,47); 2) **N**o **D**ifference (**ND**) when CLAMP occupancy between HS and NHS conditions is non-differential by EdgeR (46,47) but exhibits a <25% difference in ChIP-seq coverage; 3) **L**ow **D**ifference (**LD**) when CLAMP occupancy between HS and NHS conditions is non-differential by EdgeR (46,47) but exhibits a >=25% difference in ChIP-seq coverage. Because each loop has two ends, the loop anchors were further categorized into five types: HD-HD, HD-LD/ND, ND-ND, LD-ND, and LD-LD (**Figure S6A-F**). We found that most CLAMP-bound chromatin loop anchors **gained (Up)** (65/71: 92%) and **lost (Down)** loop anchors (849/980: 87%) upon HS belong to the ND or LD categories, in which CLAMP occupancy on chromatin is not dramatically changed. Thus, we conclude that upon HS, most CLAMP-bound chromatin changes occur through modulation of 3D chromatin contacts between loop anchors rather than through the gain or loss of CLAMP occupancy at specific genomic locations, consistent with the rapid time scale of heat stress-induced transcriptional and phase changes.

We also measured the average number of genes present: 1) at the loop anchors (**Table S7**) and 2) within the CLAMP-bound HS-induced differential chromatin loops (**Table S8**). We quantified the average loop length of CLAMP-bound loops. We found that CLAMP-bound loops were gene-rich, both at the loop anchors and within the loops, with the average CLAMP-bound HS-induced chromatin loop size smaller (avg 38 kb) compared to global HS-induced differential loop size (avg 172 kb) (**Figure 3C-D and Figure 2C-D**).

Since the majority of CLAMP-bound loops were down-regulated by heat shock, we investigated what proportion of all HS-induced Down-loops were CLAMP-bound chromatin loops and whether they overlapped with HS-repressed genes (**Figure 3E**). Although most 3D loop changes after HS are CLAMP-dependent (∼85%, Figure 2E), only ∼11% of HS-induced Down-loops were CLAMP-bound chromatin loops (**Figure 3E**), of which 122 sites overlapped with CLAMP-dependent HS-repressed genes. This suggests that CLAMP modulates 3D looping through both **direct** mechanisms that likely involve specific combinations of cofactors and **indirect** mechanisms that may involve modulation of the activity of other chromatin loop regulators in the cell.

To visualize the types of loops that CLAMP mediates directly to regulate heat shock repression, we focused on a genomic region on chromosome 2R, where we identified a subset of global 3D chromatin loop changes that are **directly** bound and regulated by CLAMP (**Figure 3F**). We show that a subset of CLAMP-bound chromatin loops are differentially affected by HS, and corresponding genes within these loops are repressed after HS. At the CLAMP-associated loops, before and after HS, there is no significant difference in CLAMP occupancy (**Figure 3G, V and VII**); however, there is differential chromatin loop formation, with a subset of chromatin loops lost while others are formed after HS (**Figure 3G, VI and VIII**). The repression of these target genes is CLAMP-dependent, as shown by the nascent RNA tracks from HS and NHS control and *clamp* RNAi-treated cells (**Figure 3G, IX, X**). Therefore, CLAMP directly represses a subset of its target genes through modulating chromatin looping.

Next, we integrated the CLAMP HiChIP data with the nascent RNA sequencing and micro-C datasets to identify CLAMP-dependent HS-repressed genes (N=735), which are: 1) **directly** CLAMP-bound and CLAMP-dependent HS-induced Down-loops (14.4%, N=106); 2) **indirectly** CLAMP-dependent HS-induced Down-loops (52.8%, N=388), and 3) independent of HS-induced loop changes (29.5%, N=217) (**Figure 4A and Table S9**). The first gene category comprises repressed genes directly regulated by CLAMP at the 3D chromatin organizational and transcriptional level after HS. In contrast, the second gene group is indirectly regulated by CLAMP at the 3D chromatin organizational level after HS. In contrast, the last group is regulated by CLAMP at only the transcriptional level and not the 3D chromatin organizational level (**Figure 4A).** Thus, repression of genes after HS due to changes in 3D chromatin architecture at ∼67% of HS-repressed genes is either directly or indirectly regulated by CLAMP. In contrast, ∼30% of the CLAMP-dependent HS-repressed genes are not associated with concurrent changes in 3D loop structure after HS, indicating a 3D loop-independent transcriptional regulation of a subset of CLAMP-dependent HS-repressed genes (**Figure 4A**). Therefore, we conclude that CLAMP directly and indirectly inhibits 3D chromatin loop formation during HS at a subset of gene-rich genomic locations, resulting in their transcriptional repression.

**Figure 4.**
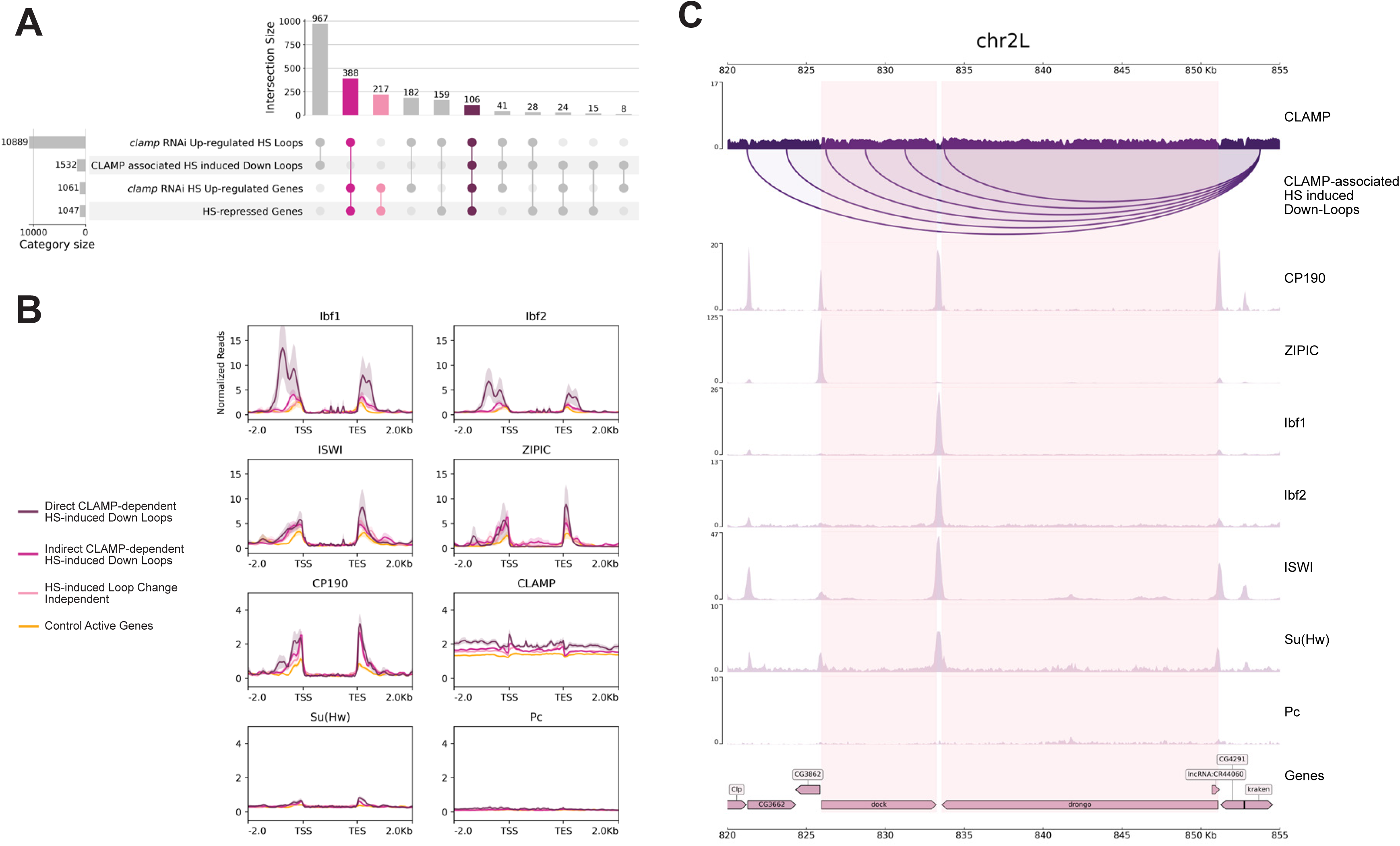
HS-associated chromatin loops directly and indirectly regulated by CLAMP have different chromatin-bound factors under NHS conditions. **A.** Upset plot comparing all CLAMP-dependent (Micro-C data, *clamp RNAi* up-regulated HS Loops) and CLAMP-bound (HiChIP data) chromatin loops lost after HS with CLAMP-dependent HS-repressed genes, showing a significant (Fisher’s Test) number of CLAMP-dependent HS-repressed genes are associated with CLAMP-dependent loss of chromatin loops after HS (p=1.55E-04). Notably, a significant proportion of these are directly CLAMP-bound (p=8.78E-32). **B.** Schematic showing three categories of CLAMP-dependent HS-repressed genes based on 3D chromatin organisational changes associated with HS. **C.** Average profile for chromatin-bound factors across CLAMP-dependent HS-repressed genes present at direct and indirect CLAMP-dependent HS-induced Down-loop anchors, genes not associated with any loop anchors, and a random set of genes (negative control). **D.** 35 kb region on 2L showing distribution of chromatin bound factors at CLAMP-dependent HS-repressed genes *dock* and *drongo* and associated CLAMP-bound chromatin loops lost after HS

### 4. Specific combinations of cofactors present prior to heat stress distinguish the direct from the indirect functions of CLAMP in 3D looping and heat stress repression

Next, we asked the question: What combination of chromatin-associated factors distinguishes CLAMP-dependent repressed genes that are directly versus indirectly regulated by CLAMP? To address this question, we screened publicly available architectural protein datasets (GSE11804, GSE30740, GSE36374, GSE36393, GSE54529, GSE63518, GSE80702, GSE89244) (12,26–33), which were previously used to examine co-factors for Pipsqueak (also, a GA-binding transcription factor similar to CLAMP) and Polycomb in 3D chromatin organization (26). We measured the occupancy of known chromatin marks and binding factors prior to HS at the CLAMP-dependent HS-repressed genes (N=735), which are associated with: 1) direct CLAMP-dependent HS-induced Down-loops (N=106); 2) indirect CLAMP-dependent HS-induced Down-loops (N=388); and 3) HS-induced loop change independent (N=217) (**Figure 4A, B, and Table S9**).

We identified factors present before HS at the CLAMP-dependent genes associated with CLAMP-bound chromatin loops (**direct** regulation) and those associated with CLAMP-independent chromatin loops (**indirect** regulation) to identify candidate co-factors involved in directly regulating HS-induced transcription repression by CLAMP. We used a set of random genes not repressed after HS as a control (**Figure 4B**). Genes in all three categories lacked repressive factors such as Su(Hw) and Pc (26,48) (**Figure 4B**). Compared with genes that are not repressed by CLAMP, the CLAMP-dependent HS-repressed genes (N=735) (both **direct** and **indirect** effects) were enriched for: 1) ISWI, a chromatin remodeler (53) at their TSS and TES (transcription start and end) sites which is known to require CLAMP for its recruitment (36); 2) CP190, a BTB-ZnF factor present in most insulator complexes; and 3) ZIPIC (Zinc-finger protein interacting with CP190), a promoter-bound TF with architectural and insulator function (49–52) (**Figure 4B**). Interestingly, genes where CLAMP has a **direct** effect on repression through regulating 3D loops (N=106) were more enriched for the CP190-interacting insulator binding factors: Ibf1 and Ibf2 (49) (**Figure 4B**). Thus, we hypothesize that Ibf1 and Ibf2 are important cofactors that regulate the direct function of CLAMP in repression of genes through 3D looping.

The presence of a chromatin remodeler like ISWI and insulator proteins like ZIPIC and CP190 prior to HS at genes that will be repressed may prime these genes for subsequent transcription repression. Upon HS, CLAMP is **directly** required for the loss of the chromatin 3D loop structure and repression at a subset of these genes that are enriched for Ibf1/2 (**Figure 4C**). In contrast, genes enriched for ZIPIC and CP190, but lacking Ibf1/2 and CLAMP at the chromatin loops, still undergo 3D chromatin loop loss and repression after HS, but CLAMP is **indirectly** responsible for their 3D chromatin loop changes. Future work is required to identify the factors that regulate the CLAMP-dependent **indirect** changes in 3D chromatin organization and transcriptional repression: ZIPIC and CP190 are strong candidate factors to regulate indirect effects. Also, CLAMP-dependent repressed genes (**Figure 4B**) that are not associated with any 3D chromatin changes do not show enrichment for ISWI or insulator proteins compared to genomic regions undergoing 3D changes. How CLAMP regulates repression independent of 3D looping requires further investigation and may involve modulating Pol II pausing because we have shown that CLAMP directly interacts with the NELF pausing regulator (17).

Overall, our studies have identified CLAMP as the first reported DNA-binding factor that regulates HS-dependent repression and identified a mechanism that involves changes in 3D chromatin loops. Moreover, we provide the highest resolution data set for studying 3D chromatin loops after heat stress in *Drosophila*. Our data are consistent with a model in which CLAMP-regulated inhibition of chromatin loop formation after HS is responsible for repression of genes after HS, suggesting that these loops would normally be important for maintaining active transcription (**Figure 5**). We also identify Ibf1/2 as candidate cofactors that are enriched when CLAMP functions **directly** versus **indirectly** in regulating transcriptional repression through 3D looping. Future work will be required to define the combinatorial context-specific interactions between CLAMP, chromatin remodelers, and insulators that determine how genes become repressed upon heat stress.

**Figure 5.**
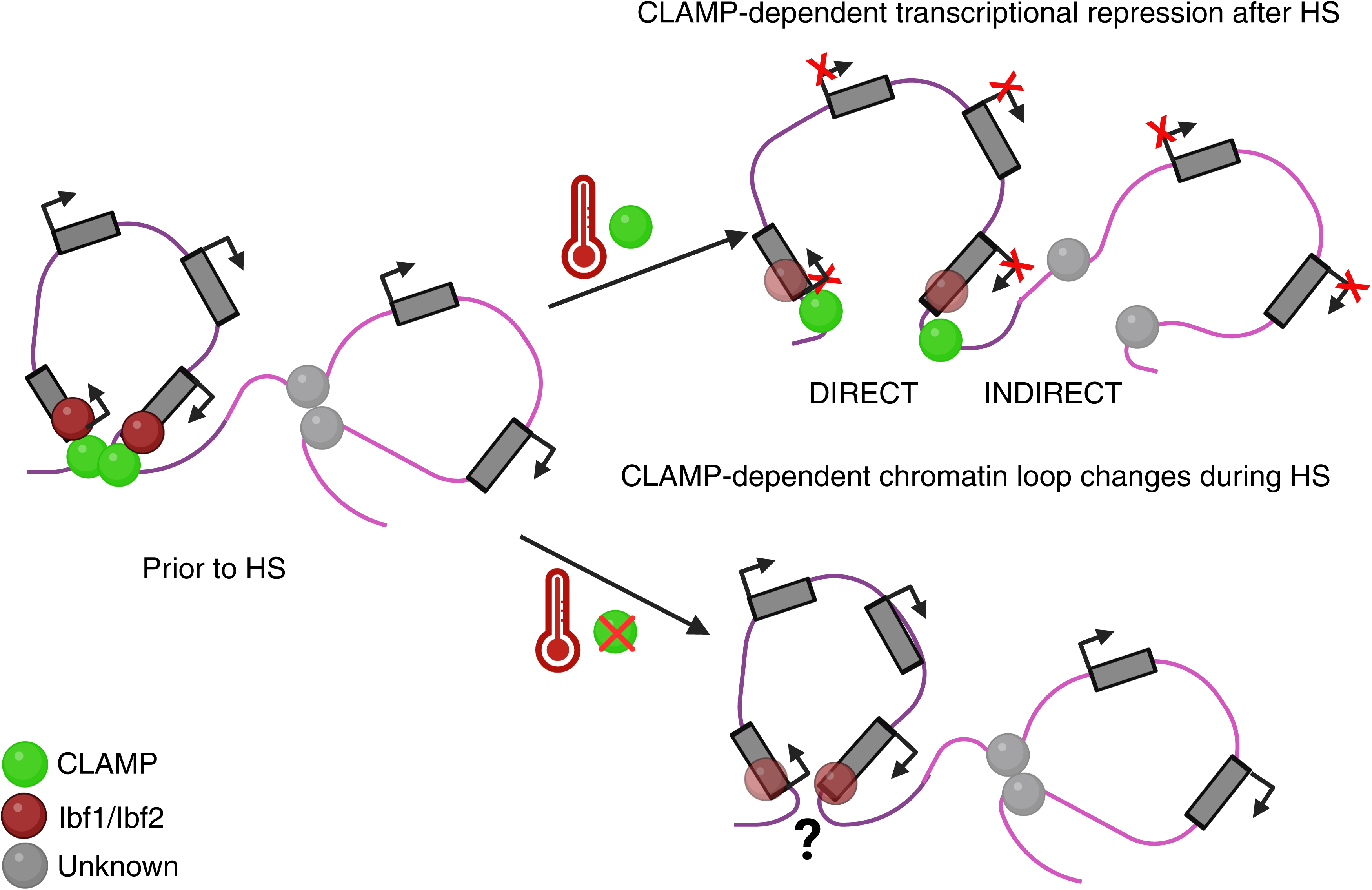
Model for CLAMP-dependent HS-induced transcriptional gene repression.

## Discussion

Temperature is one important abiotic regulator of gene expression that substantially regulates biological functions. Defining how increasing temperature affects gene transcription has not only identified new transcriptional mechanisms but is vital for managing the effects of global changes in temperature on living organisms, including humans, plants, and other animals.

To survive under heat stress, cells must repress the ongoing transcription of constitutive gene transcripts to avoid accumulating toxic protein products, which would lead to cell death. Compared with decades of work on transcriptional activation upon heat stress (3–5,7), very few reports have provided insight into how HS-induced transcriptional repression is regulated (3,7,10–12,54). In particular, sequence-specific DNA-binding TFs that target genes for transcriptional repression remained unknown.

Here, we demonstrate that the GA-binding transcription factor CLAMP is the first-reported TF that controls the repression of the majority of genes (**Figure 1**) and genome-wide rearrangement of 3D chromatin loops (**Figure 2E**) upon heat stress. In contrast, CLAMP does not regulate any of the canonical genes activated upon heat stress, such as the heat shock protein genes (**Figure 1, Table S2a-d**). A significant number of CLAMP-dependent HS-repressed genes overlap with CLAMP-dependent 3D loop changes (**Figure 2H-I, Table S1**), indicating the role of CLAMP in regulating HS-dependent transcription repression via 3D chromatin structure changes. However, the loss of CLAMP-associated loop anchors alone is insufficient to determine whether a gene will be activated or repressed because CLAMP-associated loop anchors are lost upon HS at both repressed and activated genes (**Figure 2**).

Furthermore, we found that only a subset of CLAMP-dependent repressed genes are associated with the CLAMP-bound 3D chromatin loops that are lost upon heat stress (**Figure 4A**), suggesting a more restricted direct and a broader indirect role for CLAMP in regulating local changes in 3D genome organization for HS-induced transcription repression. However, CLAMP is the first report of any DNA-binding factor involved in HS-induced changes in gene repression and chromatin organization across species. Also, direct mechanisms of transcriptional dysregulation are always a subset of all transcriptional changes caused by the loss of a transcription factor *in vivo,* and therefore we asked whether there are specific cofactors enriched prior to heat stress at genes where CLAMP functions **directly** compared with sites where it functions **indirectly**.

On average, CLAMP-dependent HS-repressed genes are modestly enriched prior to heat stress for ZIPIC, a zinc finger protein (**Figure 4B**). ZIPIC is an architectural protein with homodimerization domains that promotes CP190-mediated insulator activity and boundary formation, which can prevent transcription of nearby genes (50–52). CP190 is another protein modestly enriched at the CLAMP-dependent HS-repressed genes. CLAMP physically interacts with CP190 and both promote gypsy chromatin insulator activity (55). Additionally, the enrichment of ISWI, an essential chromatin remodeler that can be recruited by CLAMP (17) at HS-repressed genes compared to control active genes, further supports the hypothesis that a change in chromatin architecture drives HS-induced gene repression. Also, most (67%) CLAMP-dependent HS-repressed genes are associated with HS-dependent chromatin loop changes (N=388+106) regulated by CLAMP, either **directly** or **indirectly** (**Figure 4A, Table S1**).

We found that CLAMP-dependent HS-repressed genes, at which CLAMP **directly** regulates chromatin looping, were enriched for the Ibf1/2 insulator proteins that interact with CP190 (49) (**Figure 4, 5**). However, future work is necessary to define the CLAMP-interacting factors at loci where CLAMP indirectly regulates 3D chromatin changes during HS without binding to the loop anchors, as well as at the smaller group of loci where HS-induced transcriptional repression is loop-independent. Furthermore, how the combinatorial context-specific interactions between all the factors we identified and other unknown factors determine whether a gene will become repressed upon heat stress still needs to be understood.

Also, we have not yet determined how HS results in 3D chromatin changes at CLAMP-bound loops, especially in places where there is no concurrent loss in CLAMP occupancy after HS. We predict that loss of the loop after HS would result in transcriptional repression due to loss of interactions that activate transcription through 3D looping or 2D mechanisms such as RNA Polymerase II pause release (7). In addition, we show that the direct effect of CLAMP on chromatin looping to mediate repression is associated with binding prior to HS of the insulator chromatin factors (Ibf1/2) and not the Polycomb and Su(Hw) complexes. However, it is possible that after heat stress, the occupancy of these factors changes, and the biomolecular mechanisms by which CLAMP and Ibf 1/2 drive 3D chromatin structural changes remain unclear.

CLAMP is an intrinsically disordered protein with a prion-like domain, PrLD (13,40). Therefore, we speculate that heat stress may result in biophysical changes at CLAMP-associated loop anchors that regulate the liquid-liquid phase separation properties of bound proteins. We have recently shown that CLAMP itself can undergo a phase transition (40). Several reports suggest that increases in temperature influence phase separation properties of disordered proteins with prion-like domains (PrLD) (10,56–60). Therefore, it is possible that HS induces changes in the phase separation properties of CLAMP at the loop anchors. Also, CLAMP interacts with Negative Elongation Factor (NELF) that forms condensates under stress to drive transcriptional repression and regulates Pol II pausing (10,17). In the future, it will be interesting to define how the CLAMP-NELF interaction and the relationship between CLAMP andIbf1/2 regulates HS-induced transcriptional repression. The presence of a different combination of chromatin-bound factors at different CLAMP-bound target genes may regulate the phase transition properties of the locus, which could determine whether a gene will be activated or repressed upon HS. However, further experiments need to be done to test this hypothesis using *in vitro* chromatin models.

Overall, we show that the GA-binding TF CLAMP is a key DNA-binding factor that promotes transcriptional repression after HS more strongly than activation. Insight into transcriptional repression is important because the mechanism of HS-induced transcriptional repression has remained much less understood than HS-induced activation across species. We also generate new high-resolution 3D and gene expression data sets that can be used for developing combinatorial predictive models of how gene expression is rapidly modulated under different conditions. However, future work is necessary to define the combinatorial context-specific interactions between all of the factors we identified and other unknown factors that determine whether a gene will become activated or repressed upon heat stress.

## Author contribution

M.R. and E.N.L. planned experiments, analyzed results, and wrote the manuscript. J.L.A., M.R., and K.C. performed micro-C experiments. M.R. and J.L.A. performed the HiChIP experiments. M.R. and J.D. performed the SLAM-seq experiments. J.D. performed computational analysis of the SLAM-seq data. J.L.A. and A.A. performed computational analysis of micro-C and HiChIP data, generated and integrated both data sets with available ChIP data sets, and generated SLAM-seq data sets. J.L.A. and J.D. reviewed and edited the manuscript. S.M.L. performed the polytene chromosome squashes and immunostaining after HS.

## Conflict of interest

The authors declare no conflicting interests.

## Funding

This work was supported by R35GM126994 to E.N.L. from NIH (National Institutes of Health (NIH). J.L.A. is funded by the HHMI Gilliam Scholar Program. S.M.L. was funded by Brown University’s Undergraduate Teaching and Research Awards (UTRAs). K.C. is a Blavatnik scholar.

## Notes

### Competing Interest Statement

The authors have declared no competing interest.

### Summary of Updates

Figures 2, 3, and 4 updated, and changes made to the main title. Changes to the text were made to clarify the discussion. Added New Supplementary Figure S1.

