## Supplementary Figures and Legends for "The CLAMP GA-binding transcription factor regulates heat stress-induced transcriptional repression"

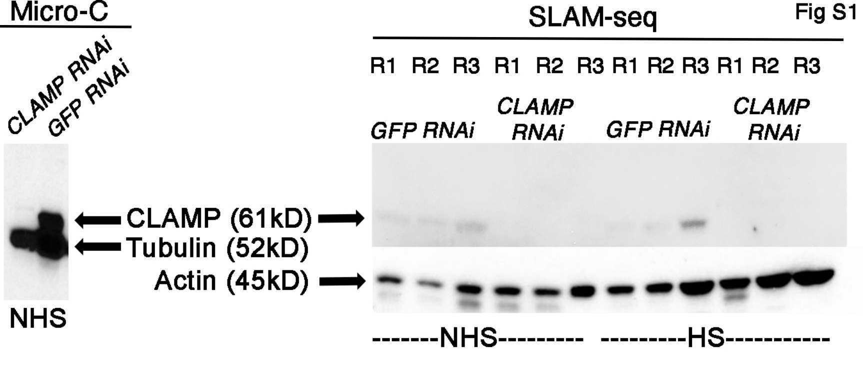

Fig S1
Micro-C
CLAMP RNAi
GFP RNAi
NHS
CLAMP (61kD)
Tubulin (52kD)
Actin (45kD)
SLAM-seq
R1 R2 R3 R1 R2 R3 R1 R2 R3 R1 R2 R3
GFP RNAi
CLAMP RNAi
GFP RNAi
CLAMP RNAi
-------NHS---------
---------HS-----------

Fig S2

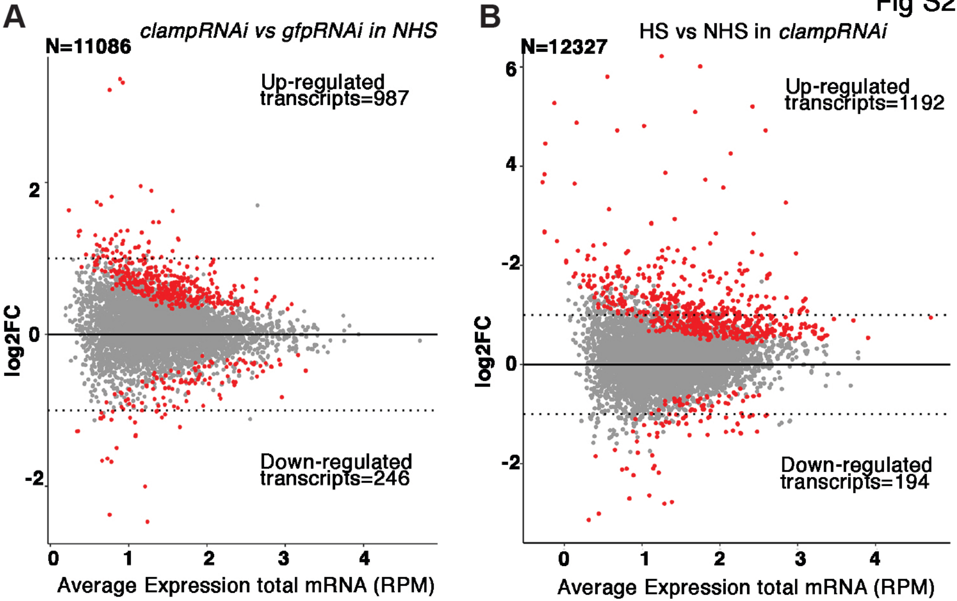
A
clampRNAi vs gfpRNAi in NHS
N=11086
Up-regulated transcripts=987
Down-regulated transcripts=246
log2FC
2
0
-2
0
1
2
3
4
Average Expression total mRNA (RPM)
B
HS vs NHS in clampRNAi
N=12327
Up-regulated transcripts=1192
Down-regulated transcripts=194
log2FC
6
4
-2
0
-2
0
1
2
3
4
Average Expression total mRNA (RPM)

Fig S3

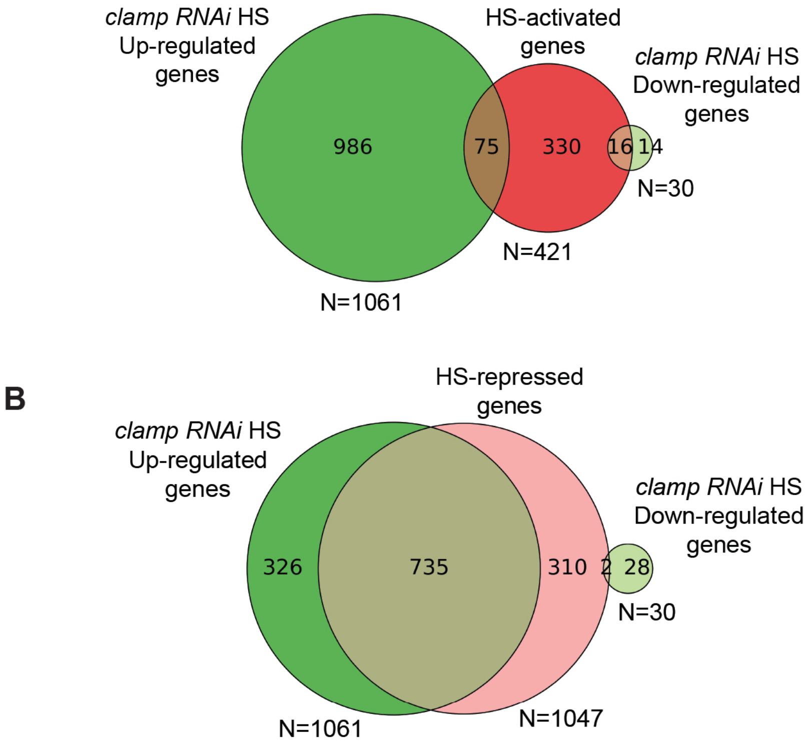
A
clamp RNAi HS Up-regulated genes
HS-activated genes
clamp RNAi HS Down-regulated genes
986
75
330
16
14
N=30
N=421
N=1061
B
HS-repressed genes
clamp RNAi HS Up-regulated genes
clamp RNAi HS Down-regulated genes
326
735
310
2
28
N=30
N=1061
N=1047

Fig S4

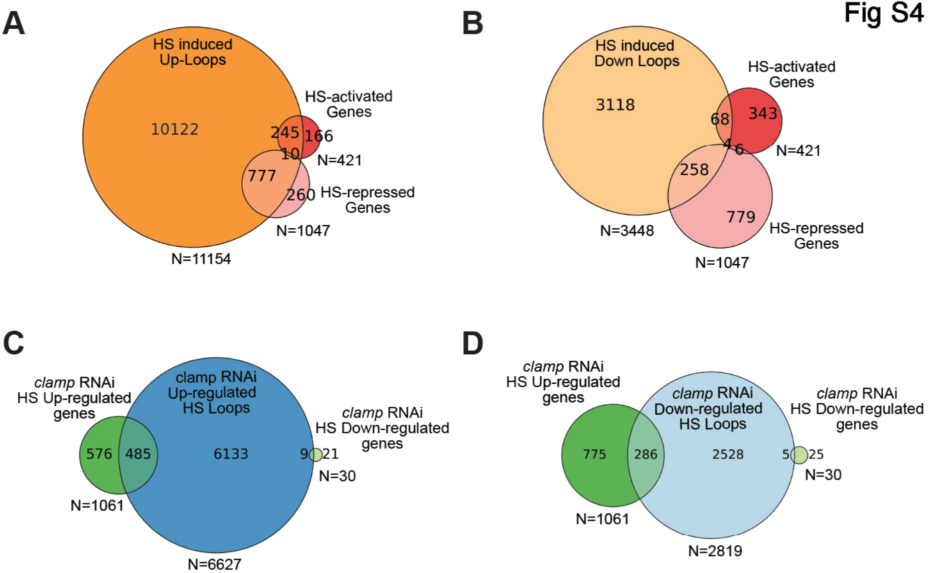
A
HS induced Up-Loops
10122
245
166
10
777
260
HS-activated Genes
N=421
HS-repressed Genes
N=1047
N=11154
B
HS induced Down Loops
3118
68
343
4
6
258
779
HS-activated Genes
N=421
HS-repressed Genes
N=3448
N=1047
C
clamp RNAi HS Up-regulated genes
clamp RNAi Up-regulated HS Loops
clamp RNAi HS Down-regulated genes
576
485
6133
9
21
N=30
N=1061
N=6627
D
clamp RNAi HS Up-regulated genes
clamp RNAi Down-regulated HS Loops
clamp RNAi HS Down-regulated genes
775
286
2528
5
25
N=30
N=1061
N=2819

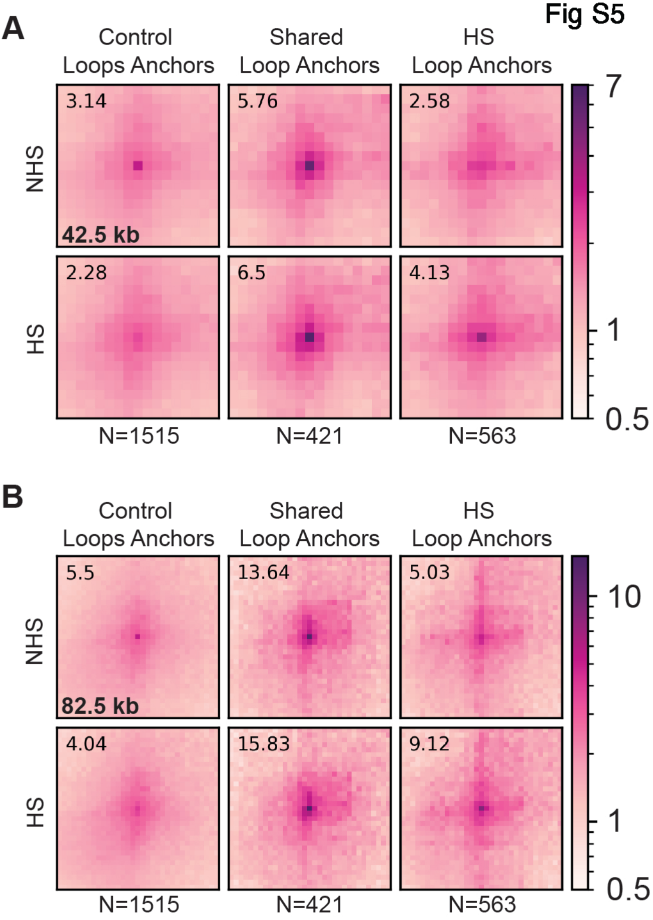
Fig S5
A
Control
Loops Anchors
Shared
Loop Anchors
HS
Loop Anchors
NHS
HS
3.14
5.76
2.58
42.5 kb
2.28
6.5
4.13
N=1515
N=421
N=563
7
1
0.5
B
Control
Loops Anchors
Shared
Loop Anchors
HS
Loop Anchors
NHS
HS
5.5
13.64
5.03
82.5 kb
4.04
15.83
9.12
N=1515
N=421
N=563
10
1
0.5

Fig S6

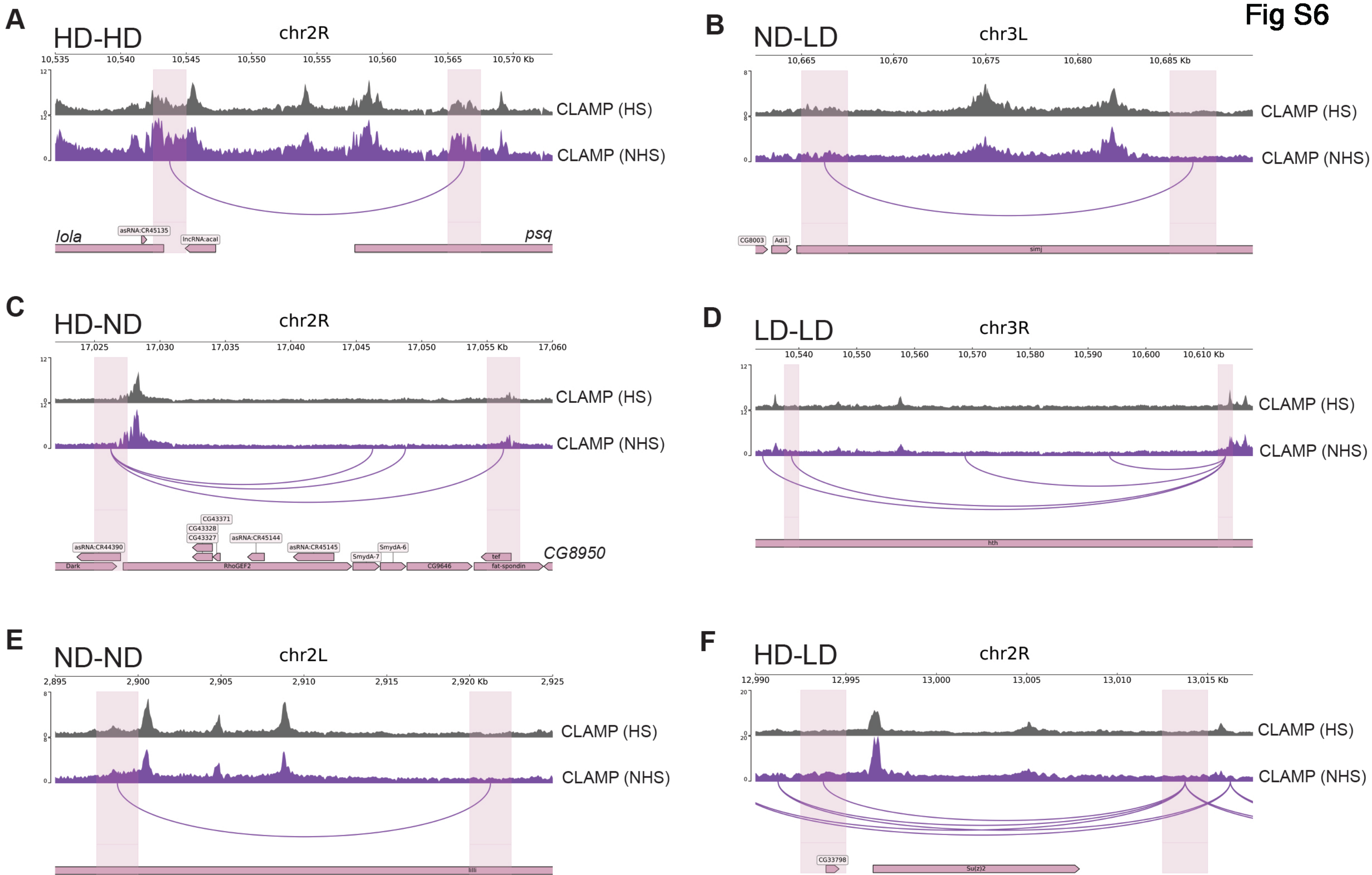
A
HD-HD
chr2R
10,535
10,540
10,545
10,550
10,555
10,560
10,565
10,570 Kb
CLAMP (HS)
CLAMP (NHS)
lola
asRNA:CR45135
lncRNA:acal
psq
B
ND-LD
chr3L
10,665
10,670
10,675
10,680
10,685 Kb
CLAMP (HS)
CLAMP (NHS)
CG8003
Adi1
simj
C
HD-ND
chr2R
17,025
17,030
17,035
17,040
17,045
17,050
17,055 Kb
17,060
CLAMP (HS)
CLAMP (NHS)
CG43371
CG43328
CG43327
asRNA:CR44390
asRNA:CR45144
asRNA:CR45145
SmydA-7
SmydA-6
tef
CG8950
Dark
RhoGEF2
CG9646
fat-spondin
D
LD-LD
chr3R
10,540
10,550
10,560
10,570
10,580
10,590
10,600
10,610 Kb
CLAMP (HS)
CLAMP (NHS)
hth
E
ND-ND
chr2L
2,895
2,900
2,905
2,910
2,915
2,920 Kb
2,925
CLAMP (HS)
CLAMP (NHS)
lilli
F
HD-LD
chr2R
12,990
12,995
13,000
13,005
13,010
13,015 Kb
CLAMP (HS)
CLAMP (NHS)
CG33798
Su(z)2

### Supplementary figure legends

**Fig. S1 Photomicrograph showing validation of CLAMP dsRNA treatment, lowering CLAMP protein level in Kc cells**

**Fig. S2 CLAMP functions more frequently as a repressor than an activator in both NHS and HS conditions.**

**A-B.** MA plots show more up-regulated nascent transcripts (positive red dots, $p < 0.05$, log2FC $\geq$ 2) in both *clamp RNAi*-treated cells under NHS (no heat stress) and HS (heat stress) conditions.

**Fig. S3 CLAMP-dependent HS-repressed genes significantly overlap with HS-repressed genes A-B.** Venn diagrams comparing HS-activated, N=421 (**A**), and HS-repressed, N=1047 (**B**) genes with *CLAMPRNAi*-dependent HS-upregulated (N=1061) and downregulated (N=30) genes. *CLAMPRNAi* upregulated genes significantly overlap with HS-repressed genes (p=0, Fisher's test with FDR correction)

**Fig. S4 Many CLAMP-dependent repressed genes are present at the loop anchors of CLAMP-dependent HS-associated chromatin loops lost after HS.**

**A-D.** Venn diagrams comparing HS-associated chromatin loop changes (genes at the loop anchors) and HS-activated and repressed genes (**A** and **B**); as well as CLAMP-dependent chromatin loop changes during HS with CLAMP-dependent HS-induced transcriptional changes (**C** and **D**), showing significant (Fisher's Test) overlap between HS-associated Up loops (**A**) and Down loops (**B**) with HS-repressed genes; and CLAMP-dependent HS-associated Down loops (**C,** *clamp RNAi* up-regulated HS-loops) and Up loops (**D,** *clamp RNAi* down-regulated HS-loops) with CLAMP-dependent HS-repressed genes (*clamp RNAi HS* up-regulated genes).

**Fig. S5 CLAMP-HiChIP identifies CLAMP-bound chromatin loops in control and HS cells.**

**A-B** APA plots of CLAMP HiChIP loops centered on three classes of significant loop interactions identified by CLAMP HiChIP in both NHS and HS conditions: 1) Loops identified only in NHS conditions (control loops, N=1515); 2) Loops identified in both NHS and HS conditions (shared loops, N=421) and 3) Loops identified only in HS conditions (HS loops, N=563). The value listed indicates the enrichment of the center pixel compared to the background, at ~40 kb (**A**) and ~80 kb (**B**) resolution.

**Fig S6. Most CLAMP-associated changes upon heat stress are three-dimensional and not associated with differences in CLAMP occupancy.**

**A-F.** Example tracks for each class of CLAMP-associated loops lost and gained after HS. The grey tracks show the occupancy of CLAMP upon HS, whereas the purple tracks show CLAMP occupancy in normal conditions (NHS). A purple line connects loop anchor endpoints. Highlighted regions correspond to 2.5 kb anchors. The classes of CLAMP-associated loops shown are: **A**) HD-HD, in which there is a change in both three-dimensional contacts and CLAMP occupancy at both ends of the loop anchor upon HS; **B**) ND-LD in which there is a change in 3D contacts and only a slight change in CLAMP occupancy at one of the ends of the loop anchor upon HS; **C**) HD-ND in which there is a change in 3D contacts and occupancy of CLAMP at one end of the loop anchor upon HS; **D**) LD-LD in which there is a change in 3D contacts and only a slight change in CLAMP occupancy at both of the ends of the loop anchor upon HS. **E**) ND-ND in which there is only a change in 3D contacts and no change in CLAMP occupancy at either end of the loop anchor upon HS; **E**) HD-LD in which there is a change in 3D contact at one end of the loop anchor and only slight change in CLAMP occupancy at another end of the loop anchor upon HS.

### Supplementary Table legends

**Table S1:** List of statistical tests and their results listed for the Venn diagrams generated for analysis of micro-C, HiChIP, and SLAM-seq data sets.

**Table S2a-d:** List of genes down and up-regulated in control (*gfp RNAi*) cells after HS compared to NHS (a), in *clamp* RNAi-treated cells compared to control (*gfp RNAi*) cells in NHS condition (b), in *clamp* RNAi-treated cells after HS compared to NHS condition (c), and in *clamp* RNAi-treated cells compared to control (*gfp RNAi*) cells under HS condition (d).

**Table S3-** List of HS-regulated and CLAMP-dependent RNA transcripts from nascent RNA-sequencing SLAM-seq data.

**Table S4**- List of HS-regulated and CLAMP-dependent genes from nascent RNA-sequencing SLAM-seq data.

**Table S5-** List of genes present at the loop anchors of the HS up and down loops and those regulated by *clamp* RNAi under HS conditions.

**Table S6-** List of genes present within the chromatin loops, HS-induced up and down loops, and those regulated by *clamp* RNAi under HS conditions.

**Table S7-** List of genes present at the loop anchors CLAMP-associated HS-induced up and down loops.

**Table S8:** List of genes present within CLAMP-associated HS-induced up and down chromatin loops.

**Table S9-** List of CLAMP-dependent repressed genes which are directly and indirectly regulated by CLAMP at the 3D chromatin structural level after HS.
